# Metatranscriptomic analysis indicates prebiotic effect of Isomalto/malto-polysaccharides on human colonic microbiota *in-vitro*

**DOI:** 10.1101/2023.12.31.573771

**Authors:** Klaudyna Borewicz, Bastian Hornung, Fangjie Gu, Pieter H. van der Zaal, Henk A. Schols, Peter J. Schaap, Hauke Smidt

**Author notes:** Corresponding author: Bastian Hornung, CBG-MEB, Graadt van Roggenweg 500, 3531AH Utrecht, Netherlands. These authors contributed equally to this work. Mead Johnson, Middenkampweg 2, 6545 CJ Nijmegen, The Netherlands. CBG-MEB, Graadt van Roggenweg 500, 3531AH Utrecht, Netherlands. TUMCREATE, 1 CREATE Way, CREATE Tower, #10-02 Singapore 138602, Singapore. IFF, Willem Einthovenstraat 4, 2342 BH, Oegstgeest, The Netherlands.

## Abstract

Isomalto/malto-polysaccharides (IMMPs) are a novel type of soluble dietary fibres with a prebiotic potential promoting growth of beneficial microbes in the gut. However, the mode of action of IMMPs remains unknown. Previous studies on IMMPs showed an increase in total bacteria, especially lactobacilli, and higher production of short chain fatty acids (SCFA) when IMMPs were fed to rats or used during *in vitro* fermentation. Here we investigated with metatranscriptomics how IMMPs with different amounts of α-(1→6) glycosidic linkages affected microbial function during incubation with human faecal inoculum. We showed that active microbial community dynamics during fermentation varied depending on the type of IMMP used and that the observed changes were reflected in the community gene expression profiles. Based on metatranscriptome analysis, members of *Bacteroides*, *Lactobacillus* and *Bifidobacterium* were the predominant degraders of IMMPs, and the increased gene expression in these bacteria correlated with high amounts of α-(1→6) glycosidic linkages. We also noted an increase in relative abundance of these bacteria and an activation of pathways involved in SCFA synthesis. Our findings could provide a baseline for more targeted approaches in designing prebiotics for specific bacteria and to achieve more controlled modulation of microbial activity towards desired health outcomes.

## Introduction

The human gut is home to a diverse ecosystem inhabited by bacteria, archaea, viruses and eukaryotes, which play an important role in their host’s health and well-being ^1–3^. These organisms interact with each other and with the host via a complex network of relationships, and knowing the mechanisms of these interactions and how to influence them might provide a useful tool for refining the function of this ecosystem to promote homeostasis and to strengthen host’s immunity against infections ^4^. Currently ways to manipulate the composition and function of gut microbiota range from mild measures, such as use of dietary supplements (especially pro- and prebiotics ^5^) or implementation of dietary regimes, to more extreme measures, such as the use of antibiotics ^6^ or faecal transplantations ^7^. Prebiotics are complex carbohydrates, often soluble dietary fibres, that cannot be digested by human enzymes but are readily used by the colonic microbiota and provide a health benefit for the host ^8^. A range of different prebiotics may stimulate growth and activity of specific microbial groups (e.g. butyrogenic bacteria ^9^), leading to the production of different metabolites with health- supporting effects. However, the exact mode of action and the specific impact on microbial interactions needs to be investigated.

Isomalto/malto-polysaccharides (IMMPs) comprise a novel class of soluble dietary fibres with prebiotic potential. IMMPs are synthetized from starch by enzymatic conversion of α-(1→4) glycosidic linkages into α-(1→6) glycosidic linkages by 4,6-α-glucanotransferase (GTFB) from *Lactobacillus reuteri* 121 ^10^. The resulting α-(1→6) linkages make IMMPs resistant to digestion by human enzymes in the small intestine. As such, IMMPs can pass into the large intestine where they are fermented by the resident microbes capable of breaking down the α-(1→6) glycosidic linkages. This property of IMMPs makes them interesting as a prebiotic food ingredient. A previous *in vitro* study has reported an increased production of short chain fatty acids (SCFA), especially acetate and propionate, when IMMPs were used as a carbon source for fermentation with human faecal inoculum as the microbial source ^10^. Here we investigated the effects of three different IMMPs on microbial composition and function during *in vitro* batch fermentations with faecal inoculum from healthy human adults. Additionally, we used a fourth substrate, an IMMP preparation after removal of remaining digestible starch segments consisting mainly of α-(1→4) glucose moieties, to investigate the direct impact of having mainly α-(1→6) glycosidic linkages available to the gut microbiota.

We showed that specific changes of the microbiota, such as growth of *Bifidobacterium* and *Lactobacillus* could be attributed to the IMMPs, and that these changes were also reflected at the transcriptomic level, i.e. upregulation of specific gene groups, as well as in enzymatic activity and increase in production of SCFA.

## Results

We performed two *in vitro* batch fermentation experiments to understand how the IMMPs containing different amounts of α-(1→6) glycosidic linkages were broken down by human faecal microorganisms over time, and how the chemical structure of these compounds affected the functional dynamics of the microbial community during fermentation. Experiment A included fermentation of IMMPs of varying percentage of α-(1→6) glycosidic linkages (27%, IMMP-27; 94%, IMMP-94) at three different time points. This was complemented by experiment B that was performed with IMMP with 96% α-(1→6) linkages (IMMP-96) and IMMP-27 after treatment with α-amylase and amyloglucosidase (IMMP- dig27). Furthermore, in experiment B an additional set of time points was evaluated to provide a more detailed understanding of microbial community dynamics. In both experiments a control blank without any IMMP substrate was included. We then performed metatranscriptome sequencing of all samples and assembled the resulting data into one reference metatranscriptome. Afterwards, machine learning techniques were applied to identify groups of similarly behaving bacteria and to identify consistent dynamic patterns in gene expression.

### Quality control and statistics

The metatranscriptomes were sequenced and subjected to a quality control before the data was further analysed (Figure S1). As a result, 320 million reads (89% of the raw reads and 54% of all bases) passed the quality check and were used for contig assembly. In experiment A, the assembly yielded over 140,000 contigs, with more than 200,000 protein coding genes, and contained, on average, 81% of the input reads (range 71% - 85%) per sample. Read counts for experiment B were acquired by mapping to the same assembly obtained from experiment A (Table S1), and showed the same average mapping rate (81%, range 71% - 89%). After mapping, the biological replicates within each experiment showed a spearman correlation of on average 0.86 (range 0.78 - 0.93), indicating good reproducibility within sets of samples from the same treatment group (Table S2). The six samples from time point 0 showed a correlation of at least 0.79 (max 0.86), and all samples taken from batches incubated in the presence of prebiotics showed a correlation of at least 0.7 for time point 24h, and all (except two) showed a correlation of at least 0.6 at time point 48h, indicating similar development over all cultures.

Of the ∼200.000 protein coding genes, ∼36.000 were full length genes. To ∼144.000 protein sequences at least one Interpro domain (excluding “Coils” domain) could be identified. These included more than 24.000 genes with at least one (partial) EC number.

### Community structure and expression patterns

Taxonomic classification to at least the superkingdom of Bacteria was assigned to 190,000 of the 200,000 genes obtained from the RNA-assembly. Less than 3,000 genes were assigned to eukaryotes and less than 2,000 to Archaea. Of the bacterial groups, most genes were assigned to the orders Bacteroidales (>67,000), Clostridiales (>40,000), Lactobacillales (27,000) and Enterobacteriales (>14,000). The genus with the highest number of assigned genes was the genus *Bacteroides* (>54.000). Figure 1 shows the relative abundance of transcripts per assigned genus.

**Figure 1.**
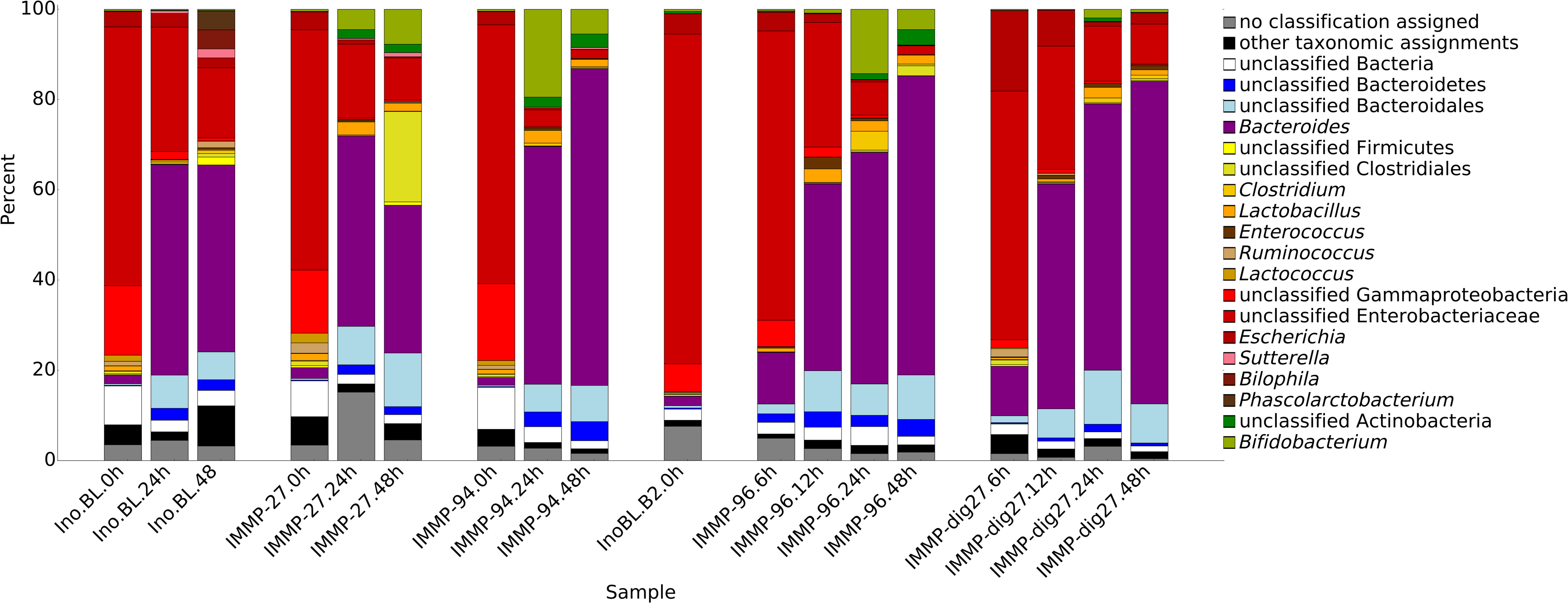
Average relative transcript expression of different genus level taxa in incubations sampled at time points 0 h, 6 h, 12 h, 24 h and 48 h. When the taxonomic assignment could not be made at genus level, the lowest classifiable taxonomy assignment was used for display. Low abundance genera are summarized as “Other taxonomic assignments” for display purposes.

To identify bacterial gene expression patterns, we focused on RNA reads for which KEGG Orthology (KO) or EC identifiers could be assigned. The percentage of reads with defined KO or EC ranged from 42% to 83% for different samples. Most of the data with assigned KO or EC identifiers came from 22 bacterial groups, of which 12 could be assigned to a known genus, and only a small number of genes was assigned to minor groups (3%), unclassifiable sequences (3%), and sequences not classifiable beyond the superkingdom Bacteria (3.5%). In the activated inoculum at the start of the incubation (t0), unclassified Enterobacteriaceae were the most active group (Figure 1), probably due to residual oxygen during the activation. However, once the incubation had started, the relative expression of *Bacteroides* increased in all treatment groups. In all samples combined across all treatments and time points, 39% of all expression data came from the genus *Bacteroides* and 27% from unclassified Enterobacteriaceae. Overall, the relative abundance of different bacterial groups based on the metatranscriptome data corresponded to the pattern in the relative abundance of different taxa based on the 16S rRNA gene analysis described previously by Gu *et al*. ^11^ (Figure 2).

**Figure 2.**
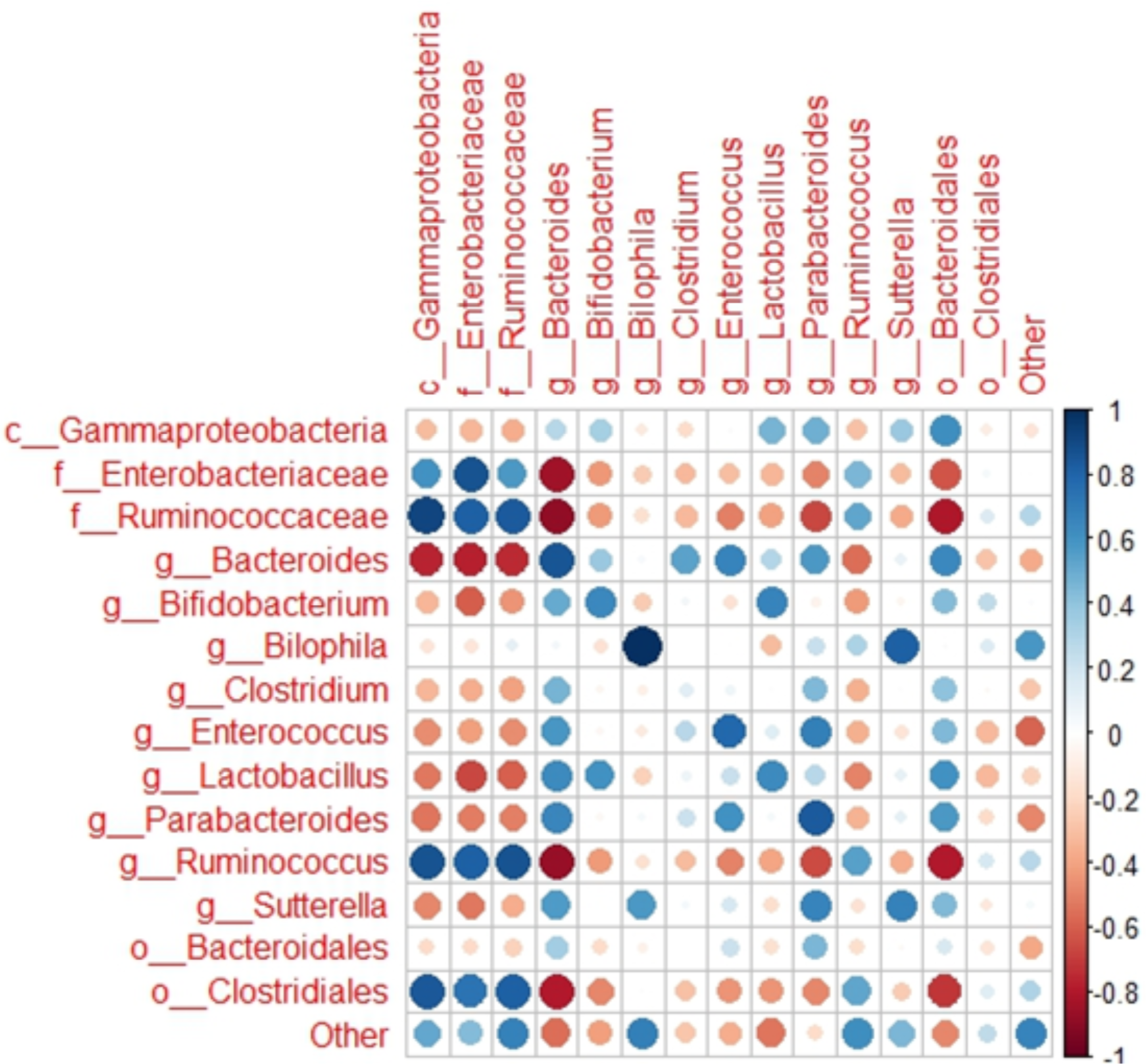
Correlation between the relative activity of the main bacterial groups based on metatranscriptome data, and their relative abundance based on 16S rRNA gene sequencing data ^11^. In case when genus level assignment was ambiguous, unclassified fraction within the next higher taxonomic level was used.

### Global and IMMP specific co-occurrence of taxa

It is known that in microbial ecosystems bacterial taxa can occupy different niches and co-exist forming a complex network of co-dependencies. We wanted to assess whether, based on the metatranscriptome data, we could identify bacterial groups which co-occurred in our samples, with particular attention to the effect of specific IMMPs. We performed clustering analysis based on mRNA reads from all samples in our dataset to test for global co-occurrence patterns. We showed that clustering into nine groups was most stable, as it could be reproduced in multiple rounds of clustering. These resulting nine different groups showed different behaviours over all investigated conditions. An overview of organism assignment per cluster, with number of assigned genes and differentially expressed genes is provided in Table S3. One of these clusters was present in all t0 samples but decreased or was absent at all other time points. This cluster consisted mostly of reads assigned to *Ruminococcus* and *Lactococcus* as well as reads that could be largely classified as contamination from the inoculum/preparation (e.g. *Homo, Mus, Bos*, unclassified Mammalia). The second and third cluster consisted mainly of genera including many genera known to contain many probiotic organism candidates, i.e. *Bifidobacterium, Lactobacillus*, and *Enterococcus,* and sequences, which could not be classified beyond a related higher taxon (e.g. unclassified Bifidobacteriaceae, unclassified Lactobacillaceae). These clusters also contained a related phage group (*Myoviridae,* mainly *Lactobacillus* phages), and an unrelated genus (*Fusobacterium*). The identified genera in cluster two and three showed increasing relative transcript abundance in all cultures supplied with IMMP substrates, whereas relative transcript abundance was decreased or undetected in the control cultures. The fourth cluster was predominated by *E. coli* and related higher order classifications (e.g. unclassified Enterobacteriaceae), together with other enterobacteria such as *Enterobacter, Citrobacter* and *Klebsiella,* and the unrelated Gram-positive genus *Eubacterium*. This cluster was mainly present in the samples without IMMP and declined in the samples with IMMP. The fifth cluster was predominated by *Bacteroides* and showed an increase with time in all incubations. This cluster also included *Parabacteroides, Prevotella, Flavobacterium*, and *Desulfosporosinus*. The sixth cluster consisted only of *Clostridium*/unclassified Clostridia, which showed some increase with time in all incubations. The seventh cluster contained *Anaerostipes* and related higher level taxon classifications (unclassified Clostridiales, unclassified Lachnospiraceae) and showed a similar pattern as cluster six. No clear pattern was seen for the eighth cluster consisting of *Corynebacterium, Ethanoligenes, Odoribacter*, and *Sutterella*. Finally, the ninth group consisted of different bacterial genera, some related to non-carbohydrate metabolizing bacteria (*Acidaminococcus, Bilophila, Phascolarctobacterium*), and some known gut symbionts like *Veillonella* and *Megasphaera*. This group was common in samples of incubations without any prebiotics at 48 h and was nearly absent in all the other samples.

### Detection of specific gene expression patterns

Besides the co-occurrence of bacterial groups, the specific gene expression patterns within these groups were investigated based on the optimal gene clustering for all bacterial groups using DBSCAN. The clustering with the optimal tau was chosen for all bacterial groups, except for the genus *Enterococcus*, for which a suboptimal tau led to better cluster separation. As a result, the DBSCAN gene clustering analysis revealed the presence of three main patterns in the expression in nearly all observed bacterial groups (Figure S3). These three patterns comprised in all cases at least 80% of all investigated genes, which were not considered noise. The first pattern was present in all incubations and was characterized by genes which were expressed only at t0, and not expressed at any of the later time points. The second pattern was found only in the control group and only at 48 h. The third and most common pattern found in all experimental groups included genes that were not expressed at t0 but showed upregulation at the later time points during incubation. This pattern was characteristic for genes assigned to the genera *Enterococcus* and *Bacteroides*, for which respectively 40% and 99% of differentially expressed genes were increasingly expressed over time in all treatment groups including the control group. *Bifidobacterium/Lactobacillus* and *Clostridium* also showed the same pattern, but only in the groups where IMMPs were present. *Anaerobutyricum hallii* (formerly *Eubacterium hallii*) showed the same gene expression pattern, but only in the group supplemented with IMMP-27 (Figure S3).

Considering the experimental set-up chosen for this study, the expression levels of genes assigned to a specific bacterial group indicate the contribution of this group to the utilization of the specified substrate, or its by-products. The high overall relative expression of genes assigned to bifidobacteria (and unclassified Bifidobacteriaceae), lactobacilli, enterococci, and unclassified Actinobacteria was positively associated with the presence of IMMPs (Figure 1). Conversely, the expression of genes assigned to unclassified Proteobacteria, *Prevotella*, *Sutterella*, *Acinetobacter*, *Eggerthella*, *Acidaminococcus*, *Streptococcus*, *Phascolarctobacterium*, and *Bilophila* was negatively associated with the presence of IMMPs, as compared to the control group.

### General metabolic effects of IMMP

We wanted to further investigate the expression of genes and corresponding bacterial groups associated with the fermentation of different IMMPs. Our analysis of the metabolic clusters revealed that five bacterial groups found in the faecal inoculum, namely *Bifidobacterium/Lactobacillus, Enterococcus, Bacteroides, Clostridium,* and *Anaerobutyricum hallii,* showed a considerable upregulation of general metabolic pathways like glycolysis, nucleic acid or fatty acid biosynthesis, as compared to the gene expression at t0. When we compared metabolic patterns between different bacterial groups, the groups exhibited overall different metabolic patterns. Members of the genus *Bacteroides* showed at first a unique partial upregulation of Vitamin B12 metabolism. An investigation of the cofactor requirements showed that Vitamin B12 in *Bacteroides* is essential for methionine synthase and methylmalonyl-CoA mutase, the latter of which produces methylmalonyl-CoA from succinyl-CoA (Figure 3).

**Figure 3.**
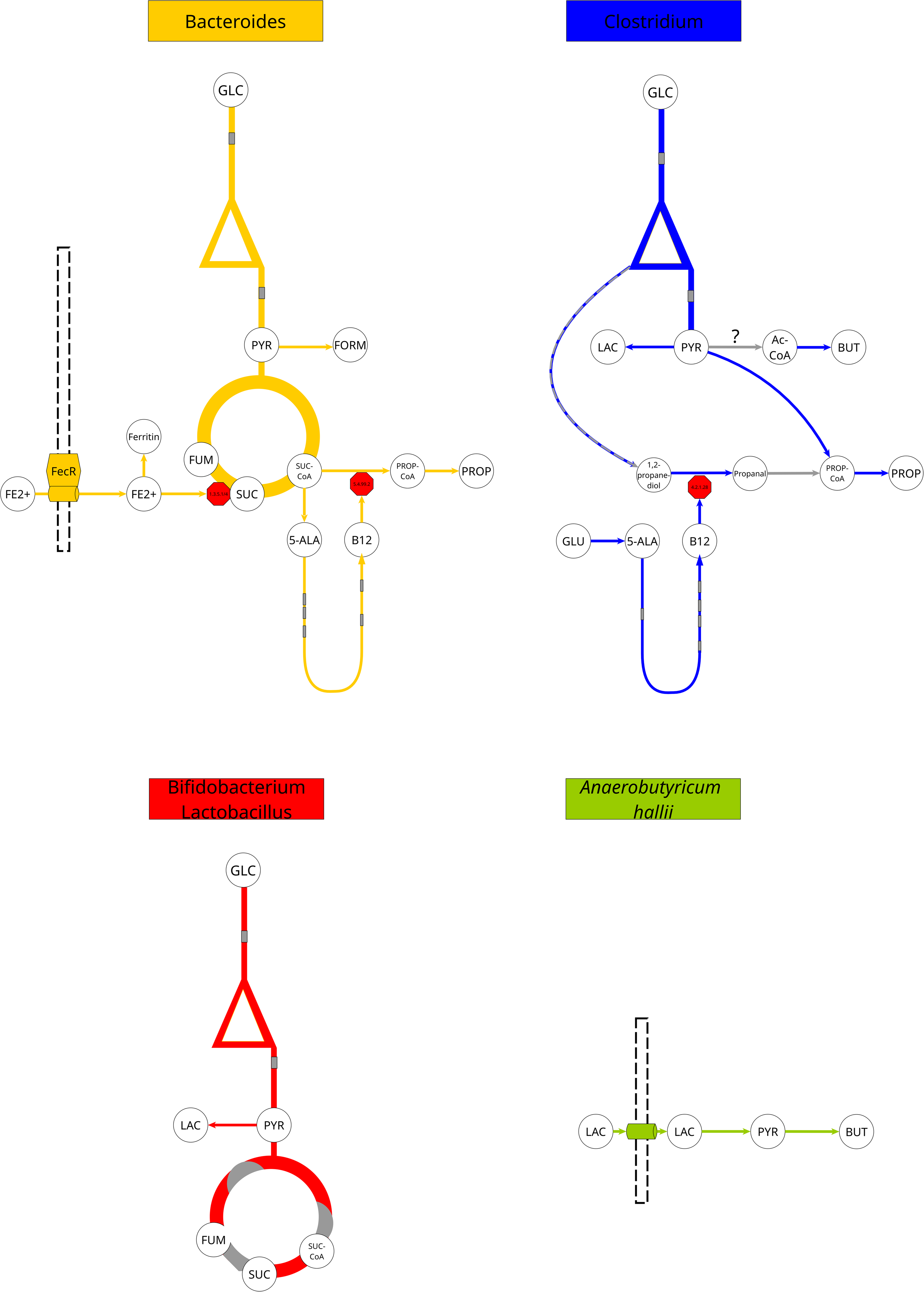
Overview of the metabolism of specific microbial groups observed in the samples taken during *in vitro* fermentation of different IMMPs by human faecal inoculum All samples show in general the same patterns for all organisms, besides for *Anaerobutyricum hallii*, which only showed expression in the samples with IMMP-dig27. The genus *Enterococcus* showed the same pattern as *Bifidobacterium/Lactobacillus*, but at lower relative transcript abundance. Grey indicates that certain genes were not differentially expressed within a pathway. 5-ALA = 5-Aminolevulinate, AC = Acetate, Ac-CoA = Acetyl- CoA, BUT = Butyrate, FORM = Formate, FUM = Fumarate, GLC = Glucose, LAC = Lactate, PROP = Propionate, PROP-CoA = Propanoyl-CoA, PYR = Pyruvate, SUC = Succinate, SUC-CoA = Succinyl-CoA

Methylmalonyl-CoA mutase is involved in propionate biosynthesis, and our data showed that the whole pathway for propionate biosynthesis was, in fact, upregulated. The data further showed that many genes coding for proteins involved in iron scavenging were also upregulated (e.g. FecR). One of the genes coding for an enzyme with iron requirements was that encoding succinate dehydrogenase, which converts succinate into fumarate. This function, as well as all others in the TCA cycle, showed upregulation in all samples tested. The genus *Clostridium* also showed an upregulation of genes involved in Vitamin B12 production, but the biosynthesis occurred via glutamate, whereas in the *Bacteroides* group it was produced via succinate. The genes in the pathway for propionate production were overall upregulated (production via acetyl-CoA, not succinyl-CoA), similar to the genes in lactate and butyrate production pathways. The only other enzyme requiring Vitamin B12 in the microbiome was a multimer of propanediol dehydratase or glycerol dehydratase (ambiguous functional assignment), both involved in the breakdown of glycerol/glycerone phosphate to propanol/propionate/1,3-propanediol. However, a full upregulation of either pathway was not observed. The *Bifidobacterium*/*Lactobacillus* group and the *Enterococcus* group showed upregulation of genes related to production of lactate from pyruvate, and the *Clostridium* group also showed upregulation of genes encoding proteins involved in butyrate production, but it is unclear if butyrate would be directly produced from pyruvate, or derived from external acetate. *Anaerobutyricum hallii*, on the other hand, showed high gene expression related to converting lactate into butyrate, as also shown previously ^12^. In addition, our data indicated that formate might have been produced by proteins encoded by genes belonging to the *Enterococcus* and *Bacteroides* populations.

### Microbial groups directly involved in the degradation of the IMMPs

In order to gain insight into which bacterial groups were directly involved in degradation of different IMMPs, we used the KEGG reference pathway for starch and sucrose metabolism ^13^. We surveyed our data for the expression of the genes encoding enzymes that are known to be involved in sucrose and starch metabolism. More specifically we focused on genes encoding enzymes from glycoside hydrolase family 13 (http://www.cazy.org/GH13_bacteria.html), as this family includes a number of bacterial proteins shown to be essential in degradation of similar compounds, such as isomaltooligosaccharides (IMOs) ^14^. The majority of genes listed in the KEGG starch and sucrose metabolism pathway were detected in our transcriptome data, as well as some additional genes in glycoside hydrolase family 13 (EC 3.2.1.135, 3.2.1.68 and 3.2.1.11), which were not listed in the KEGG pathway, but which have been shown to be activated during the degradation of pullulan and dextran ^15–18^. It is interesting to note that the relative contribution of these starch and sucrose metabolism genes to the total number of genes from each sample did not correlate with the presence or absence of IMMPs in the samples. The only exception was incubation with IMMP-dig27, in which starch and sucrose metabolism genes reached an expression of 10% at 12 h and about 12% at 48 h, whereas in other groups they ranged between 4 to 5% (Figure S4). Despite of the similarities in the overall expression of the starch and sucrose metabolism genes in all samples, we could see differences in the relative abundance of genes coding for specific enzymes depending on the IMMP used, and the duration of the fermentation (Figure S5).

One of the aims of this study was to better understand the functional dynamics of the bacterial communities during IMMP degradation. Previously reported HPAEC and HPSEC analyses ^11^ showed that the degradation of IMMP-94 and IMMP-96 occurred between 12 h and 24 h of the incubation. At 24 h and 48 h we noted an increase in the expression of genes coding for enzymes that might be directly involved in the hydrolysis of α-(1→6) glycosidic linkages, namely EC 3.2.1.10 – oligo-1,6-glucosidase, EC 3.2.1.11 – dextranase, and EC 3.2.1.33 – amylo-α-1,6- glucosidase (Figure S6a,b). There was also an increase in the expression of genes coding for enzymes that can hydrolyse α-(1→4) glycosidic linkages, mainly EC 3.2.1.1 – α-amylase, EC 3.2.1.20 – α–glucosidase 4-α-glucanotransferase, and EC 2.4.1.25 – 4-α-glucanotransferase. Since IMMP-27 contains lower amounts of α-(1→6) linkages, its degradation also involves the activation of the same genes, however, the expression levels of the genes encoding enzymes which hydrolyse α-(1→6) linkages were much lower (Figure S6a,b). Bacterial groups that contributed the most expression to the primary degradation of IMMP’s α-(1→6) linkages were *Lactobacillus*, *Bifidobacterium* and *Bacteroides*, all expressing the genes encoding EC 3.2.1.10 oligo-1,6-glucosidase and EC 3.2.1.11 dextranase. On the other hand, the metatranscriptomic data suggested that α-(1→4) linkages were hydrolysed mainly by proteins encoded by *Bacteroides*, unclassified Bacteroidales, unclassified Enterobacteriaceae, *Lactobacillus* and *Bifidobacterium* via EC 3.2.1.1 alpha-amylase and EC 2.4.1.1 glycogen/amylophosphorylase (Figure S7). Based on the transcript data, genes belonging to the genera *Bifidobacterium* and *Lactobacillus* were mainly expressed in the degradation of IMMP-94 and IMMP-96 at 24 h (Figure 4, and Figure S7). These genera showed also expression during degradation of IMMP-27 and the IMMP- dig27, but their relative contributions were much lower (Figure 4, and Figure S7). The breakdown of IMMPs at 24 h and 48 h was otherwise dominated by expression from *Bacteroides*, with the exception of IMMP-dig27 at 48 h, which showed a high level of expression of genes assigned to unclassified Enterobacteriaceae. Figure 4 summarizes our model of IMMP degradation and shows the proposed specialized role of lactobacilli and bifidobacteria in hydrolysis of α-(1→6) linkages. It also reveals the important contribution of *Bacteroides* as both primary and secondary degraders of IMMPs and their by-products.

**Figure 4.**
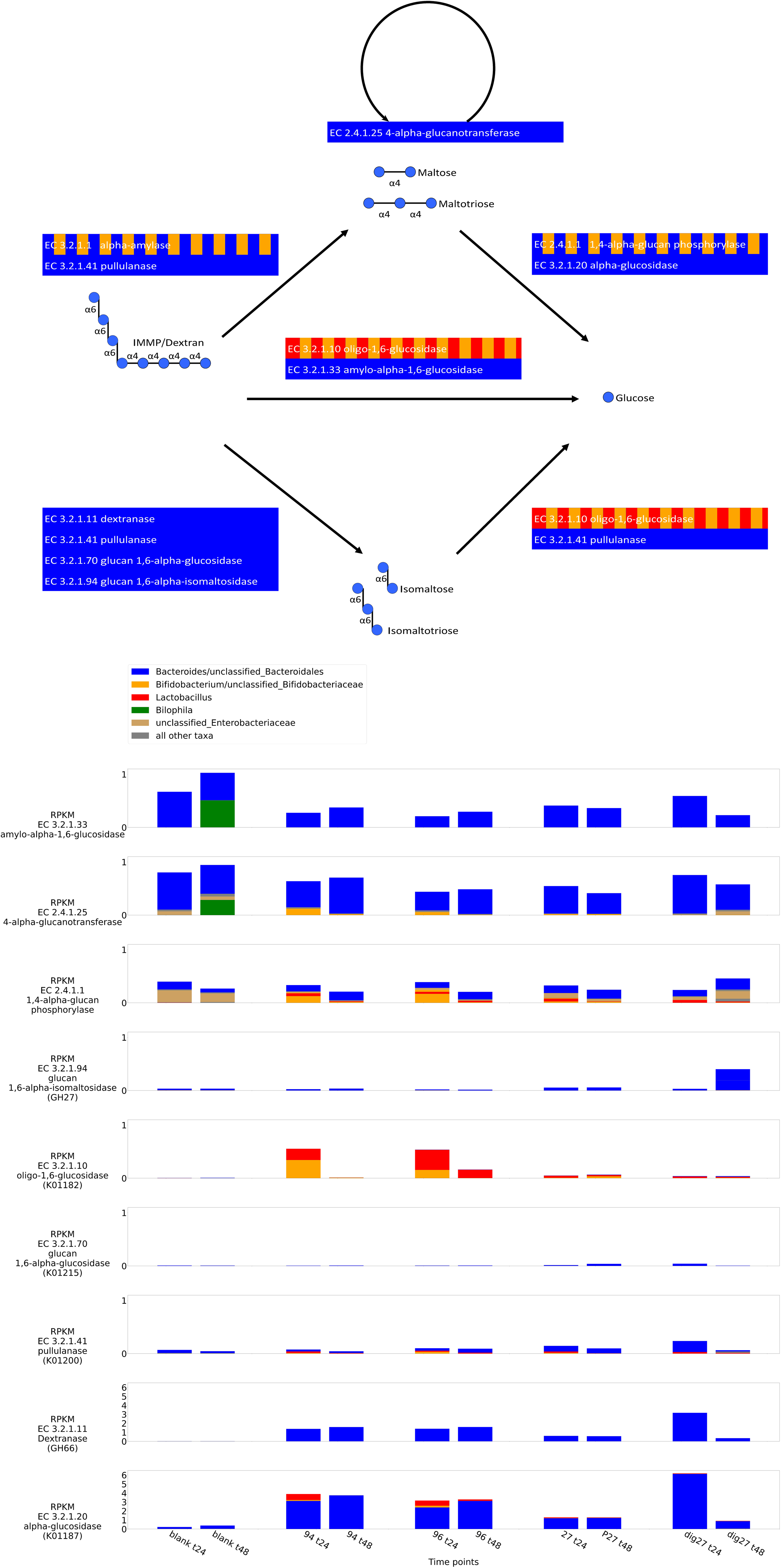
Overview over the main degradation pathways starting from dextran. Colours in the top panel indicate the main contributors to a reaction. The bottom panel shows the overall expression in reads per kilobase per million (RPKM) per organism at time points 24 and 48h for all conditions. If an enzyme could not be identified by its associated EC number, then the KO or CAZy identifier used for identification are given in brackets. Only enzymes, which were significantly differentially expressed over at least one time series, are displayed. Please note the different scales between the first and the last enzymes in the bottom panels, due to differences in expression scale.

### Comparison to other metatranscriptomic projects

We identified few limitations in our study, such as the fact that our starting culture was a mix of faecal samples from several donors for which we lack the initial gut microbiota profiles. Another limitation was the use of *in-vitro* batch fermentation system which does not allow to control for nutrient depletion and by-products accumulation. Therefore, to validate our findings we compared our results with other *in-vivo* and *in-vitro* gut metatranscriptome projects. We selected all available Illumina single-end metatranscriptomic bioprojects at the NCBI, as well as two relevant bioprojects with paired-end Illumina data. For each project we profiled taxonomic composition and compared the projects using PCA on the inverted Bray-Curtis distances. As it can be seen in Figure 5, our data showed stronger clustering than a dysbiotic microbiome project (PRJNA396964, from graft-vs-host disease patients ^19^). Our project data was also closely positioned with clusters of PRJNA416988 project ^20^, another *in- vitro* study, where samples were derived from a single person and RNA was extracted after 7 days in a bioreactor and 24h in a Hungate tube. The second most closely positioned project was PRJNA509512, where samples were derived from healthy volunteers^21^. Another *in-vitro* project, PRJNA593787^22^, with also a roughly comparable setup, was displayed further away, indicating a variation in the *in-vitro* setups comparable to *in-vivo* data.

**Figure 5.**
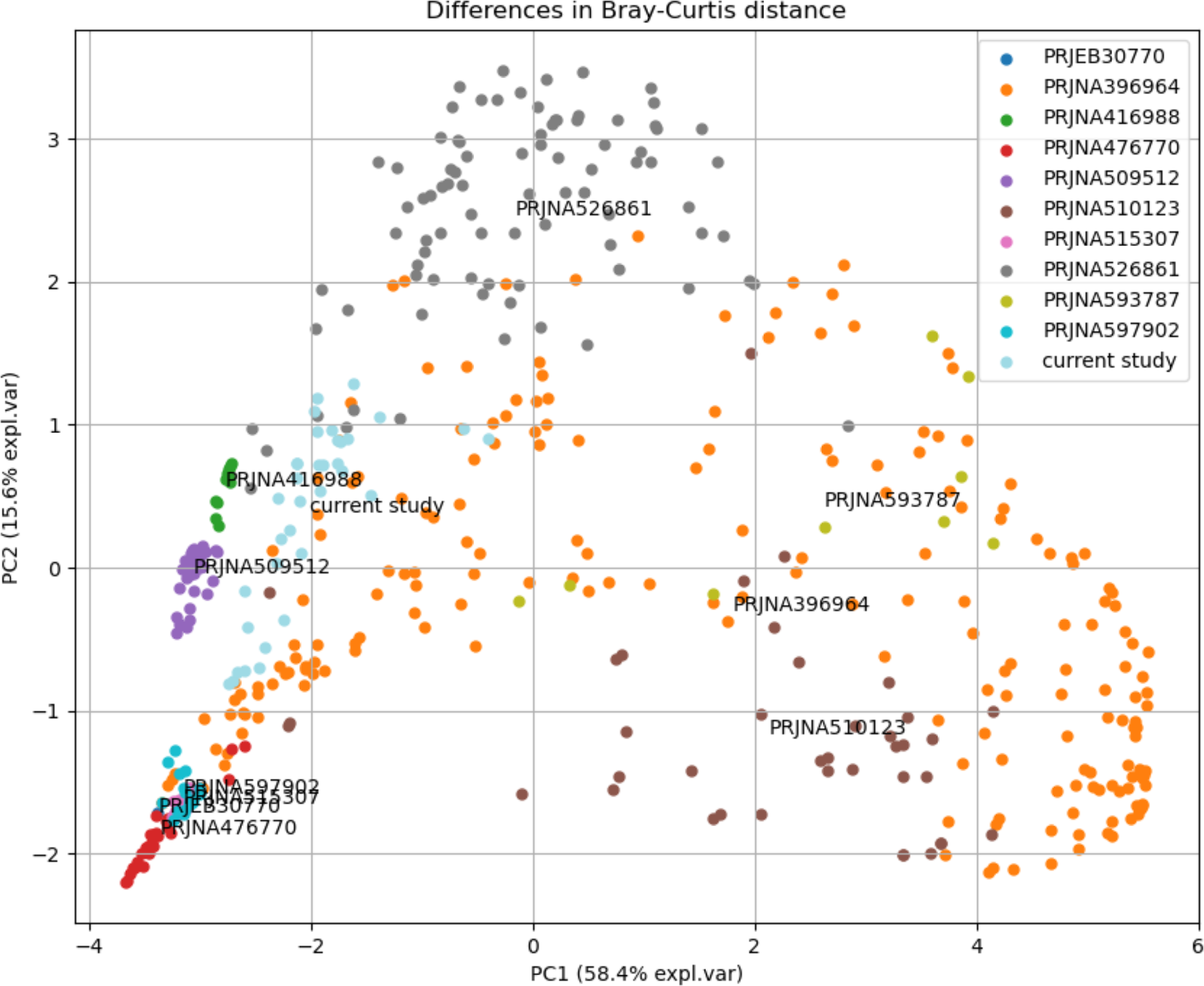
PCA based on Bray-Curtis distance between samples from different bioprojects. The current study (light blue) groups with samples from two other studies (green, purple), which are derived from either an in-vitro model based on a single volunteer or direct sequencing of healthy volunteer samples, respectively. The samples displayed in orange are derived from a graft-vs-host disease study and include disturbed microbiomes.

## Discussion

Prebiotic food components should be resistant to host’s gastric enzymes, fermentable by the host’s intestinal microbiota and capable of promoting growth and activity of bacterial groups associated with health ^8^. The IMMPs seem to fulfil all these criteria ^23–25^. Earlier studies demonstrated that (reduced) high DP IMOs are not or poorly digestible by rat gastric enzymes ^25^, and that diets containing IMOs are associated with higher numbers of lactobacilli, and an overall increase in the number of intestinal bacteria ^26^. Moreover, a recent study with human inoculum reported that IMMPs can be fermented by human large intestinal microbiota and that SCFAs, in particular acetate and propionate, are produced, indicating that IMMPs may stimulate activity of probiotic groups ^10^. This is in accordance with earlier findings from a small human trial that showed an increased level of bifidobacteria in subjects who received IMOs in their diets ^23^.

In our study, we confirmed the prebiotic character of the IMMPs and showed that the specific effect of different IMMPs on human faecal microbiota composition and gene expression varied during *in vitro* fermentation, depending on the relative amount of α-(1→6) glycosidic linkages present in the substrate (Figure 4). When IMMP-94 and IMMP-96 were used as a carbon source, we observed a strong upregulation of genes in the probiotic cluster, specifically genes assigned to bifidobacteria and lactobacilli. Furthermore, high relative gene expression of these bacteria corresponded with an increase in their relative abundance as estimated by rRNA gene sequencing ^11^. In contrast, when the IMMP-dig27 also containing mainly α-(1→6) glycosyl units was used as a substrate, the relative expression of genes assigned to bifidobacteria and lactobacilli was lower. These bacteria showed lower expression in the control group, and their highest gene expression in the presence of IMMP-27 was delayed to 48 h. Interestingly, all IMMP treatment groups showed a time lag between the maximum relative expression, and the increase in the corresponding bacterial relative abundance as measured by rRNA gene-targeted community analysis ^11^. For example, the maximum expression of bifidobacterial genes was observed at 24 h of incubation when IMMP-94 and IMMP-96 were used as substrates. Yet, bifidobacteria reached their highest relative abundance only at 48 h when relative expression of their genes had already decreased. The relative gene expression of lactobacilli followed a pattern similar to that of bifidobacteria in all treatment groups, except for incubations with IMMP-94 where genes assigned to lactobacilli showed maximum relative expression at 12 h, whereas bifidobacterial gene expression peaked at 24 h. Relative expression of genes assigned to *Bacteroides* was very high in all groups, regardless of the incubation time, or presence and type of the IMMP used. *Bacteroides* spp. are known to be generalists that are able to break down a wide array of carbon sources ^27^. Bifidobacteria and lactobacilli are often more specialized and can grow on substrates that are not necessarily accessible to other bacteria in the microbial ecosystem (e.g. human milk oligosaccharides ^28, 29^, galactooligosaccharides ^30^, other complex carbohydrates^31^) . This may be the reason that these groups show delayed gene expression in relation to *Bacteroides*, as only after the depletion of the easily accessible IMMP fractions containing the α-(1→4) glycosidic linkages the bacterial groups capable of utilizing the α-(1→6) glycosidic linkages gained a competitive advantage.

While relative abundance and gene expression of some of the beneficial bacteria increased with the presence of IMMPs, we also noted that the exclusive use of these prebiotics put a selective pressure on other beneficial microbes. For example, *Lactococcus lactis* and *Ruminococcus bromii* - two specialized beneficial degraders, did not show any survival in our samples ^11^. This can be explained by the lack of suitable substrate for both species, given that neither any simple mono- or disaccharides (for *Lactococcus* ^32^) nor type II or III resistant starch (for *Ruminococcus* ^33^) were present in this experiment. Both organisms can be considered beneficial, but were not stimulated in the particular prebiotic environment tested here, enforcing the notion that prebiotics can selectively stimulate gene expression and growth of specific groups, whereas in general, a diverse diet may be necessary to comprehensively support a stable community of commensal microbes. An alternative explanation, which just recently got discovered, is that at least *L. lactis* could be a contamination from the medium, which we can as well not rule out ^34^, but this has not yet been shown for *R. bromii*.

In our study we also observed a clear effect of having no carbohydrate source in the control samples. With the absence of the prebiotics, there was a switch of the community from processing carbohydrates to utilizing amino acids ^35^, as indicated by the increase of relative abundance of *Acidaminococcus* ^11^. In addition, there was an increase of *Bilophila* in the control samples, which is an organism previously associated with gut dysbiosis ^36^, and which previously has been shown to decrease during the use of other prebiotics ^37^.

A total of 130 families of glycoside hydrolases, 22 families of polysaccharide lyases, and 16 families of carbohydrate esterases have been described, and many of these enzymes are encoded only by the genomes of microbes (www.cazy.org) ^38^. We surveyed our data for the presence of genes encoding the enzymes that are known to be involved in sucrose and starch metabolism, mostly genes from glycoside hydrolase family 13. Few studies up to date looked at the genetics and enzymology of degradation of IMMPs mainly in lactobacilli ^16, 39, 40^, bifidobacteria ^41^ and *Bacteroides*. However, microbial species in the gut do not act in isolation, but rather can interact with each other through a network of syntrophic interactions often making the utilization of the substrate more effective ^42^. Metabolic potential and fermentation efficiency vary between different species, and complete degradation of IMMPs in the gut is a result of different bacterial groups working together in a complementary fashion, likely leading to the formation of microbial food chains ^42, 43^. Certain bacterial groups may show a higher expression of specific genes coding for enzymes required to catalyze given reactions at specific degradation steps. This was also visible in our experiments. The expression of oligo-1-6-glucosidase encoding genes was dominated by lactobacilli and bifidobacteria when IMMP-94 and IMMP-96 were used as a substrate, whereas *Bacteroides* and unclassified Bacteroidales showed also high gene expression with IMMP-27 or IMMP- dig27 substrate. Similar patterns could be observed in expression of other genes that code for enzymes involved in sucrose and starch metabolism (Suppl. Figure S7). While some of the carbohydrate breakdown steps were dominated by gene expression of known probiotic genera, many of the primary and secondary degradation processes were also performed by members of *Bacteroides*. Our data showed that once the fermentation started, one of the very specialized enzymes, dextranase, was expressed only by *Bacteroides*. Other processes were found reliant on multiple genera as based on the gene expression data. For example, the breakdown of IMMP27 and IMMP-dig27 to maltose and maltotriose by α–amylases was dominated by gene expression of the genus *Bacteroides*, whereas further metabolisation was performed also by bifidobacteria and lactobacilli. Furthermore, other groups such as enterobacteria or *Parabacteroides* were not involved in most of these breakdown processes, but still constituted viable populations in the communities. Their functional role in the community is, however, not clear.

Previously published experimental results of this experiment showed that the administration of IMMPs lead to an increased production of different SCFAs, mainly acetate and succinate ^11^. While succinate normally does not accumulate in this medium ^44^, the excess of substrate ^45^, high CO_2_ levels, and the upregulation of all the necessary steps ^46^ in our metabolic mapping, including the necessity for iron, could explain such accumulation. In addition, previous studies showed that succinate accumulation is associated with oversupply of complex substrates ^47^, such as prebiotics, or in our case IMMPs or when further metabolization of succinate is unnecessary ^44^. It is also possible that the lack of Vitamin B12, which is necessary for propionate production ^46^, resulted in the accumulation of succinate instead of propionate. We also indeed observed an upregulation of genes involved in Vitamin B12 production. However, we were unable to unequivocally determine the exact reason based on our data.

A further limitation of our study is that we do not have the gut microbiota profile of the initial samples available, that our starting culture was a mix of seven different faecal samples, and that the overall setup of this type of *in-vitro* experiments didn’t allow for continious monitoring and controlling. We performed pH measurements ^11^, but the pH dropped as a result of SCFA production, which might have impacted the composition of the microbiota. In order to better understand the validity of our findings and how our sequencing results compare with other metatranscriptomic data from other projects (despite that differing sequencing runs have the biggest impact on the sequencing results), we compared our data with other in-vivo and *in-vitro* human metatranscriptomic studies. The results showed that our data is comparable to other metatranscriptomic projects (Figure 5), and is closely positioned to data derived from healthy volunteers and from other *in-vitro* projects. And while the exact composition of our *in-vitro* does not directly reflect the human gut (e.g. loss of many Firmicutes), we could still identify many of the known human gut microbes (Figure 1). We therefore consider this experiment as being in general similar to relevant physiological conditions.

In conclusion, dietary fibres, including modified starches such as IMMPs offer a promising, non-invasive way to intentionally manipulate gut microbiota composition.

Investigations of whole bacterial communities and understanding of the mechanisms by which microorganisms interact to degrade different dietary carbohydrates are essential for our ability to manipulate gut microbiota to benefit our health. We showed how IMMPs can increase the relative abundance and gene expression of beneficial bacteria, making these novel prebiotics potentially useful in improving host’s health from the aspect of nutrition, to achieve prevention or even alleviation of diseases.

## Materials and Methods

### *In vitro* fermentation; design and sampling

The faecal inoculum stock was prepared at TNO (Zeist, The Netherlands) from fresh faeces of seven healthy adult donors, to reduce variation within the inoculum ^48^. The stock was mixed, aliquoted and stored anaerobically at −80 °C ^48^. Sterile 20 mL anaerobic serum bottles were filled with 10 mL of the Standard Ileal Efflux Medium (SIEM; Tritium Microbiology, Eindhoven, The Netherlands). The SIEM was prepared according to Rösch et *al.* ^49^, but omitting the carbon source and Tween 80. The modified SIEM medium contained 40% (v/v) BCO medium, 1.6% (v/v) salt solution, 0.8% (v/v) MgSO4 (50 g/L), 0.4% (v/v) cysteine hydrochloride (40 g/L), 0.08% (v/v) vitamin solution and 10% (v/v) MES buffer (1 M, pH 6.0) in water. Before inoculation, a faeces stock aliquot was mixed with SIEM at 1:10 v/v and incubated overnight at 37 °C. The activated inoculum was then added to the fermentation bottles at 1% (v/v) final concentration. Three different IMMP fibres were tested, with 27% (IMMP-27), 94% (IMMP-94) and 96% (IMMP-96) of α-(1→6) glycosidic linkages as compared to the total amount of glycosidic linkages. In addition, a pre-treated IMMP-27 (IMMP-dig27) sample was included after it had been digested with α-amylase and amyloglucosidase to imitate passage through the small intestine ^11^. Samples were prepared and processed in duplicate with fibres added to individual fermentation bottles at a final concentration of 10 mg/mL. Flasks were incubated at 37 °C in an anaerobic chamber with 81% N2, 15% CO2,and 4%H2. 0.5 to 2 mL of each culture was removed at different sampling time points, depending on the experiment.

In experiment A, cultures supplied with two different prebiotic fibres (IMMP-27 and IMMP-94) and one control culture without any substrate (IMMP blank, with only the modified SIEM medium, without any carbon source) were monitored over 48 h (in duplicate), and aliquots were removed at time points 0 (up to 15 min after addition of the prebiotic), 24 h and 48 h. In experiment B, cultures were supplied with two other prebiotics, IMMP-dig27 and IMMP-96, and one culture was left with no prebiotic (IMMP blank). Experiment B was monitored for 48 hours, and samples were taken at 6 h, 12 h, 24 h and 48 h (in duplicate). An aliquot of the activated blank inoculum was taken at time point 0, just before the addition of the IMMP. All samples (18) from experiment A were subjected to metatranscriptomic sequencing. In experiment B the metatranscriptomics sequencing was done for the activated inoculum at time point t0 and for the treatment groups at all time points (17). Samples for metatranscriptomics were harvested and immediately stabilized in RNAprotect (Qiagen, Hilden, Germany) following the manufacturer’s instructions, and bacterial pellets were stored at −80 °C for up to three weeks before further processing.

### RNA extraction and Illumina sequencing

Total RNA was extracted by using the beat beating - TRIzol - column method modified from Kang et al. ^50^. Briefly, bacterial pellets were re-suspended in 100 µL TE buffer (30 mM Tris-HCl, 1 mM EDTA, pH = 8.0) containing 15 mg/mL Lysozyme, 10 U/mL of Mutanolysin and 100 µg/mL of Proteinase K. Samples were vortexed for 10 s and incubated at room temperature for 10 min, and 400 µL of RLT buffer (Qiagen, Hilden, Germany) containing 4 µL of β-mercaptoethanol was added. Samples were then vortexed, mixed with 500 µL of TRIzol Max reagent (Invitrogen, Carlsbad, CA, USA) and homogenized with 0.8 g of sterilized 0.1 mm zirconia beads for three min (3 × 1 min with cooling in between) at 5.5 ms using a bead beater (Precellys 24, Bertin Technologies). Following the beating step, samples were cooled on ice, gently mixed by inverting the tube with 200 µL of ice cold chloroform for 15 s and centrifuged for 15 min at 4 °C at 12,000 × g. The aqueous phase containing total RNA was transferred to fresh tubes and mixed with an equal volume of 70% ethanol. The mixture was placed on a Qiagen RNeasy mini column (RNeasy Mini Kit, Qiagen, Hilden, Germany) and centrifuged at 8,000 × g for 15 s to bind RNA into the column. Filtrate was discarded, and the RNA binding step was repeated until the complete sample was filtered through the column. The columns were rinsed with 350 µL of RW1 buffer (RNeasy Mini Kit, Qiagen, Hilden, Germany), and 80 µL DNAse I solution (Roche, Manheim, Germany) was applied to the column and incubated for 15 min at RT to digest DNA. The columns were rinsed twice with 350 µL RW1 buffer, and twice with 700 µL of RPE buffer (RNeasy Mini Kit, Qiagen, Hilden, Germany), following with a final wash with 80% ethanol. Columns were dried by a 2 min centrifugation at maximum speed, and total RNA was eluted with 30 µL of DNAse/RNAse free water. The total RNA concentrations were measured spectrophotometrically with an ND-1000 spectrophotometer (NanoDrop® Technologies, Wilmington, DE, USA), and residual DNA concentrations were measured with the Qubit® dsDNA BR Assay Kit (Life Technologies, Leusden, the Netherlands). Samples which contained over 10 ng/µL DNA contamination were treated with the Turbo DNAfree® Kit (Ambion, Bleiswijk, Netherlands) following manufacturer’s instructions and purified using the RNeasy Mini Kit. Total RNA quality was evaluated using the Experion RNA StdSens kit (Biorad Laboratories INC, USA), total RNA concentrations were measured with NanoDrop® and DNA contamination concentrations were measured with the Qubit®dsDNA BR Assay Kit. Between 3-5 µg of total RNA from each sample was used for mRNA enrichment with the RiboZero Bacterial rRNA Removal Kit (Illumina, San Diego, CA, USA), and the quality and quantity of enriched mRNA was assessed as described above for total RNA. Between 200- 500 ng of enriched mRNA was used for cDNA production using the ScriptSeq®v2RNA-Seq Library Preparation Kit (Epicentre, Madison, WI, USA), FailSafe®PCR Enzyme Mix (Epicentre, Madison, WI, USA) and ScriptSeq®Index PCR Primers (Epicentre, Madison, WI, USA) for amplification and barcoding of di-tagged cDNA. The PCR product presence was confirmed with gel electrophoresis using the FlashGel® System (Lonza, Rockland, ME, USA). PCR products were then purified with the HighPrep® PCR kit (MagBio Genomics, Gaithersburg, MD, USA) and concentrations of indexed cDNA were measured using the Qubit®dsDNA BR Assay Kit (Invitrogen, Carlsbad, CA, USA). Approximately 28 ng of DNA from each sample was added to a pool, and final volume of each library was adjusted to 25 µL using the HighPrep® PCR kit. Two libraries were prepared containing either 17 or 18 samples, with final concentrations of 20 ng/µL in each library. Libraries were sent for single end 150 bp Illumina HiSeq2000 sequencing (GATC, Konstanz, Germany).

### Bioinformatic processing, read assembly and annotation

The bioinformatics workflow was adapted from Davids et *al.* ^51^. SortMeRNA v1.9 ^52^ software was used to screen the metatranscriptome data against all databases deployed with the program and to remove rRNA reads. Adapters were trimmed with cutadapt v1.2.1 ^53^ using default settings. Quality trimming was performed with PRINSEQ Lite v0.20.0 ^54^ with a minimum sequence length of 40 bp and a minimum quality of 30 on both ends of the read, and as mean quality. All reads containing more than three Ns or non-IUPAC characters were discarded.

Reads from experiment A (Suppl. Figure S1) were pooled and assembled with IDBA_UD version 1.1.1 ^55^ using two rounds of assembly; firstly, with the options –min_count 200 and – min_support 5, and secondly, the reads, which could not be mapped to this assembly with bowtie2 v2.0.6 ^56^, standard parameters, were extracted, and assembled with standard options, but with the output from the previous run provided as long reads. Contigs with an A/T content of >80% were removed from the final assembly. Due to hardware restrictions, we did not include reads from experiment B in the assembly, but rather mapped reads to the assembly generated from reads obtained from experiment A as described below. Since mapping rate was comparable (table S1), it was concluded that this approach is appropriate. Prodigal v2.5 was used for prediction of protein coding DNA sequences with the option for meta samples ^57^. Protein sequences were annotated with InterProScan 5.4-47.0 ^58^ on the Dutch science grid (offered by the Dutch National Grid Initiative via SurfSara), and enriched by adding EC numbers using PRIAM version March 06, 2013 ^59^. Carbohydrate active enzymes were predicted with dbCAN release 3.0 ^60^. Further enrichment for EC numbers was obtained by matching all InterProScan derived domain names against the BRENDA database (download 13.06.13) ^61^ and using a text mining algorithm that included removal of the non-alphanumerical characters (colons, commas, brackets, etc.), partial and generic terms (type, terminal, subunit, domain, enzyme, like, etc.), as well as other smaller modifications. Details are provided in Supplementary Materials and Methods.

Read counts from experiment A and B (Figure S1) were obtained with Bowtie2 v2.0.6 ^56^ using default settings. BAM files were converted with SAMtools v0.1.18 ^62^, and gene coverage was calculated with subread version 1.4.6 ^63^. Read mappings to the RNA-assemblies were inspected with Tablet ^64^.

### Taxonomic assignments

RNA sequences from the metatranscriptome assembly were compared with Blast 2.2.29 ^65^ against the NCBI NT database (download 22.01.2014) using standard parameters, besides an E-value of 0.0001, to the human microbiome (download 08.05.2014), NCBI bacterial draft genomes (download 23.01.2014), NCBI protozoa genomes (download 08.05.2014), and the human genome (download 30.12.2013, release 08.08.2013, NCBI Homo sapiens annotation release 105). Taxonomy was estimated with a custom version of the LCA algorithm as implemented in MEGAN ^66^, but with the following changes: only hits, which exceeded a bit-score of 50 were considered, and of these, only hits with a length of more than 100 nucleotides and which did not deviate more than 10% from the longest hit were accepted.

From all sequences from the assembly that did not have a match in any of the former blast analyses, another run with the – blastn option was performed against the same databases. In case this did not yield any results, a blastp of the predicted proteins was performed against a custom version of the KEGG Orthology database (http://www.genome.jp/kegg/ko.html, download 25.04.2014). Taxonomic assignment was again performed with the LCA algorithm, and for the blastp run only hits which did not deviate by more than 10% from the hit with the maximum identity were considered.

### Differential expression

Differential expression analysis was performed separately at genus level per genus in R version 3.1.1 ^67^ with the TCC package release 1.6.5 ^68^, with 36 iterations and the combination of tmm normalization and edgeR, with an FDR=0.1. Only genes with a q-value (multitest corrected p-value) of less than 0.01 in any of the relevant comparisons were considered to be significantly differentially expressed, unless otherwise mentioned. A volcano plot can be seen in Figure S8.

### Metabolic mapping

Two rounds of clustering were performed to detect patterns in the expressed genes (Figure S2). All genera, which either had an average read count of >=10 per gene, or which exceeded 1% of all reads in any given condition, were clustered into groups based on relative counts per group using the k-means algorithm in Scipy version 1.6.1 ^69^. To determine the stability of the clustering, 50 iterations with a clustering between 1 and 20 clusters were performed, with the option “iter” set to 100.000. Afterwards the average cluster support per amount of clusters over all the iterations was computed, and additionally, the clustering was investigated with a custom python implementation of clustergrams ^70^. Within the clustered genera, genes with similar expression patterns were identified with the DBSCAN algorithm ^71^. Clustering on expression patterns was performed with ELKI 0.7.0∼20150828 ^72^, the –minpts parameter was fixed to 3 and the epsilon parameter was varied in percentages. Final clustering was evaluated using the Tau index as implemented in ELKI, and the clustering result with the best Tau was chosen, unless a lower Tau led to better cluster separation.

Only genes which were differentially expressed in at least one sampling time point in any of the incubations (i.e. Ino.BL, IMMP-27, IMMP-94, IMMP-96, IMMP-dig27), were considered in the clustering analysis. Genes were normalized per row before the clustering. All derived EC numbers were mapped with custom scripts onto the KEGG database ^13^ and visualized with Python Scipy version 1.6.1 and NumPy version 0.9.0 ^69^. Correlations were calculated with the mentioned versions of Scipy/NumPy. Differentially expressed genes were mapped separately for groups of interest, and changed functions were derived from visual inspections. Cofactor requirements were investigated with the Expasy database ^73^.

### Comparison with other projects

The NCBI was searched (June 20, 2021) for all bioprojects containing single-end Illumina sequencing data derived from the human gut microbiota, as well as relevant projects with paired-end Illumina sequencing. In total 10 projects could be identified (PRJEB30770, PRJNA396964, PRJNA416988, PRJNA476770, PRJNA509512, PRJNA510123, PRJNA515307, PRJNA526861, PRJNA593787, PRJNA597902), with in total 457 samples.

Of the 457 samples one sequencing run, SRR8567581, was excluded, due to not containing enough reads. All samples were downloaded with fasterq-dump from the sra-toolkit 2.11.0 (with option –concatenate-reads),and were classified with Kraken2 v2.1.1 with the PlusPF database, https://benlangmead.github.io/aws-indexes/k2 ^74^. Bray-Curtis dissimilarity was calculated with the beta-diversity script from KrakenTools v1.2, https://github.com/jenniferlu717/KrakenTools/. The beta-diversity measures were inverted and a PCA was performed with the pca python package, https://github.com/erdogant/pca/.

### Ethical approval and informed consent

Faeces collection in the Netherlands does not require medical ethical approval if volunteers have not undergone any form of intervention, such as in this study. However, an informed consent was signed by all donors upon providing the faecal samples.

### Data accessibility

The datasets generated and analyzed during the current study are available in the ENA repository, under accession number PRJEB13209, https://www.ebi.ac.uk/ena/data/view/PRJEB13009.

## Acknowledgements

The authors thank Jasper Koehorst (Wageningen University & Research, Laboratory of Systems and Synthetic Biology) for his help with the transcriptome annotation, and Maria Suarez-Diez and Edoardo Saccenti (Wageningen University & Research, Laboratory of Systems and Synthetic Biology) for helpful discussions. The authors want to thank Bartholomeus van den Bogert (Wageningen University & Research, Laboratory of Microbiology) for the design of Figure S1.

This research was performed in the framework of the public-private partnership CarboHealth coordinated by the Carbohydrate Competence Center (CCC, www.cccresearch.nl) and financed by participating partners and allowances of the TKI Agri&Food program, Ministry of Economic Affairs. This work was furthermore supported by Wageningen University and the Wageningen Institute for Environment and Climate Research (WIMEK) to BH through the IP/OP program Systems Biology [project KB-17-003.02-023].

This work was carried out on the Dutch national e-infrastructure with the support of SURF Foundation.

## Author contributions

KB and FG conducted the experiments. BH performed the bioinformatics processing. KB and BH analysed the data, designed figures and wrote the manuscript. HAS and HS conceived the study. PS advised on bioinformatics methods. PZ provided the IMMP substrates and provided biochemical assistance. All authors reviewed the manuscript. KB and BH contributed equally to this manuscript.

