## Supplementary figures and images for "Metatranscriptomic analysis indicates prebiotic effect of Isomalto/malto-polysaccharides on human colonic microbiota *in-vitro*"

### Supplementary figure 1

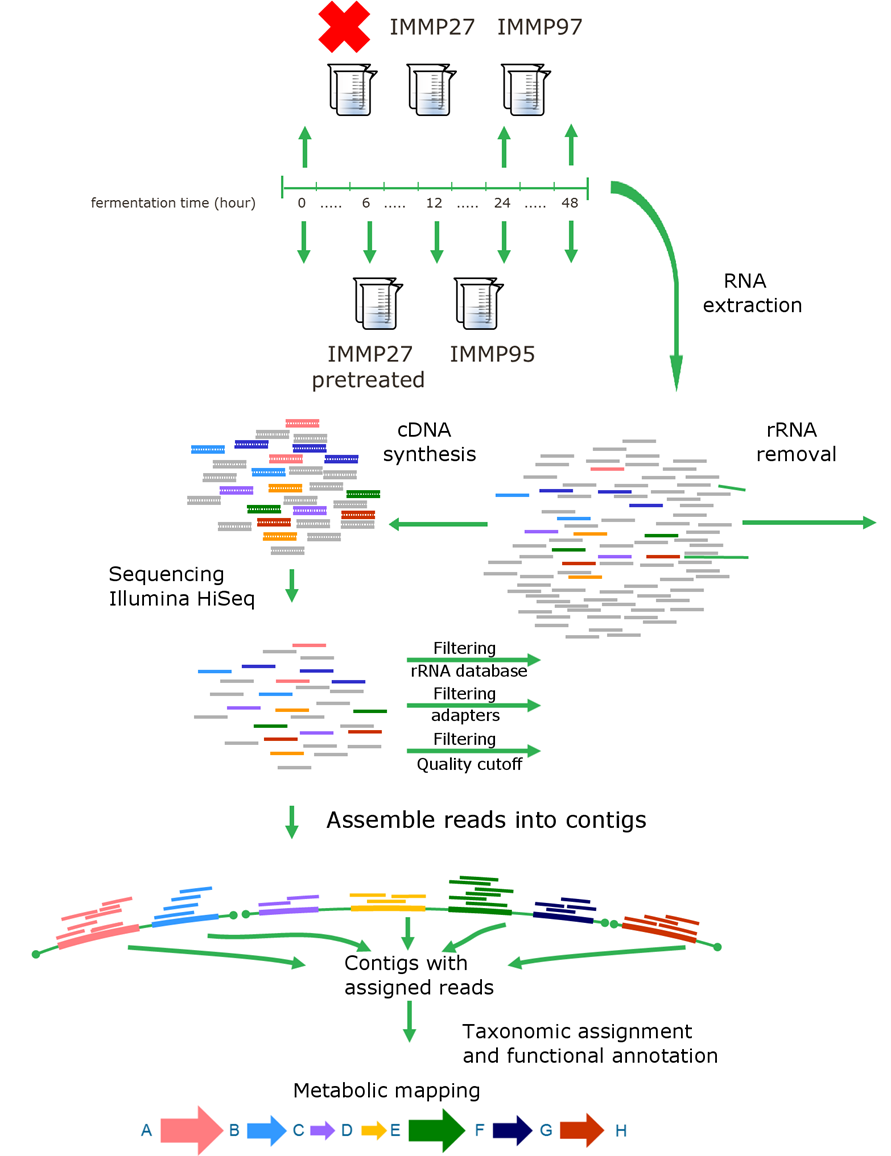

### Supplementary figure 2

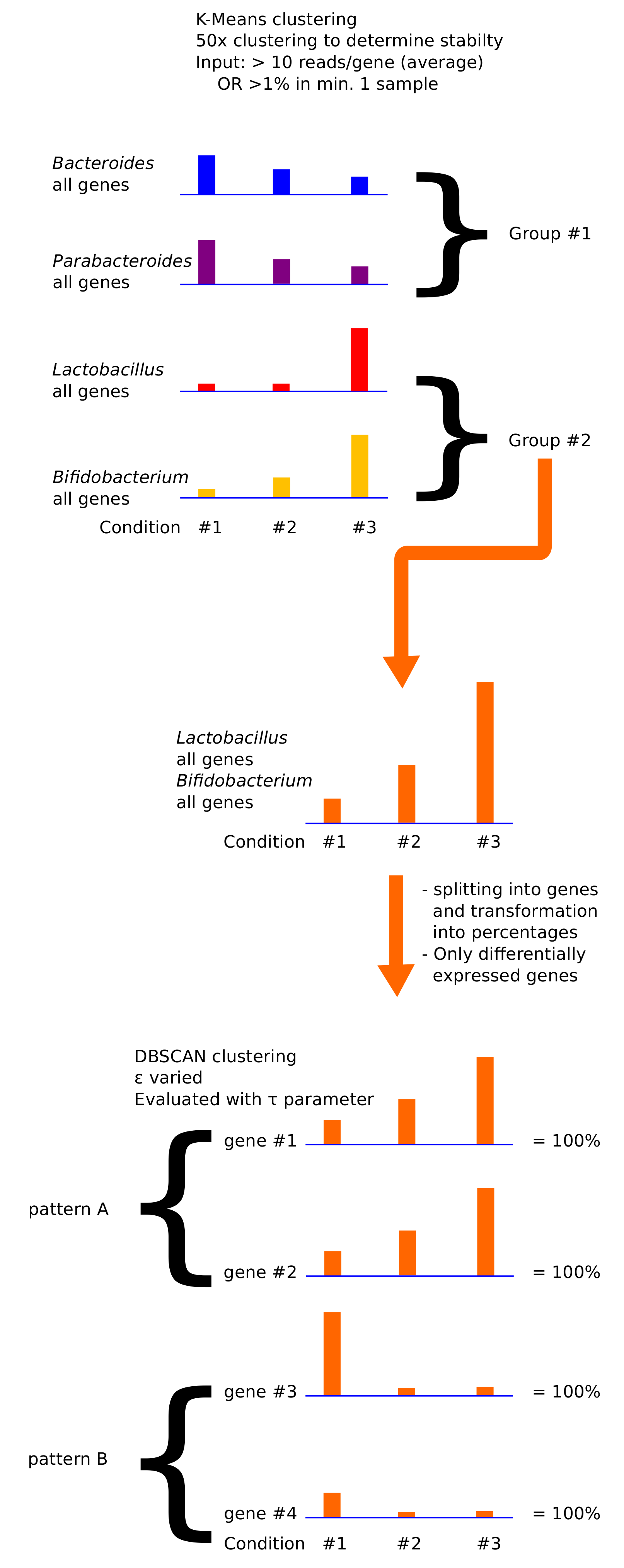

### Supplementary figure 3

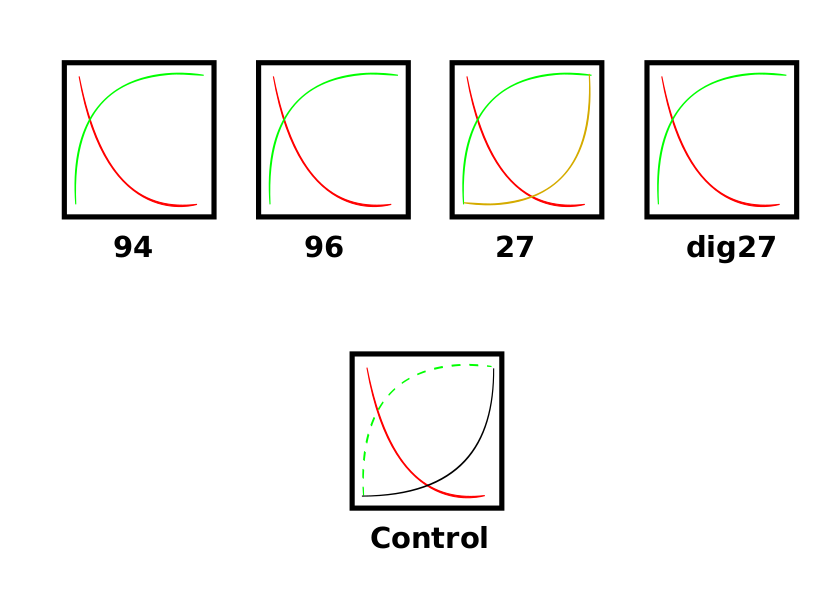

### Supplementary figure 4

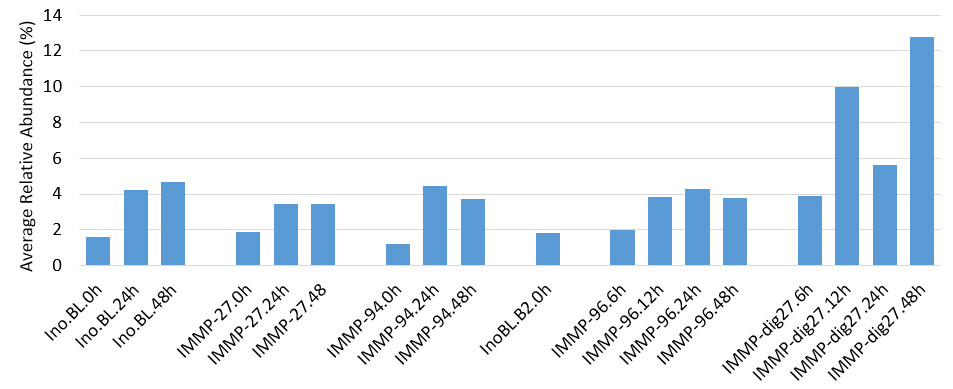

### Supplementary figure 5

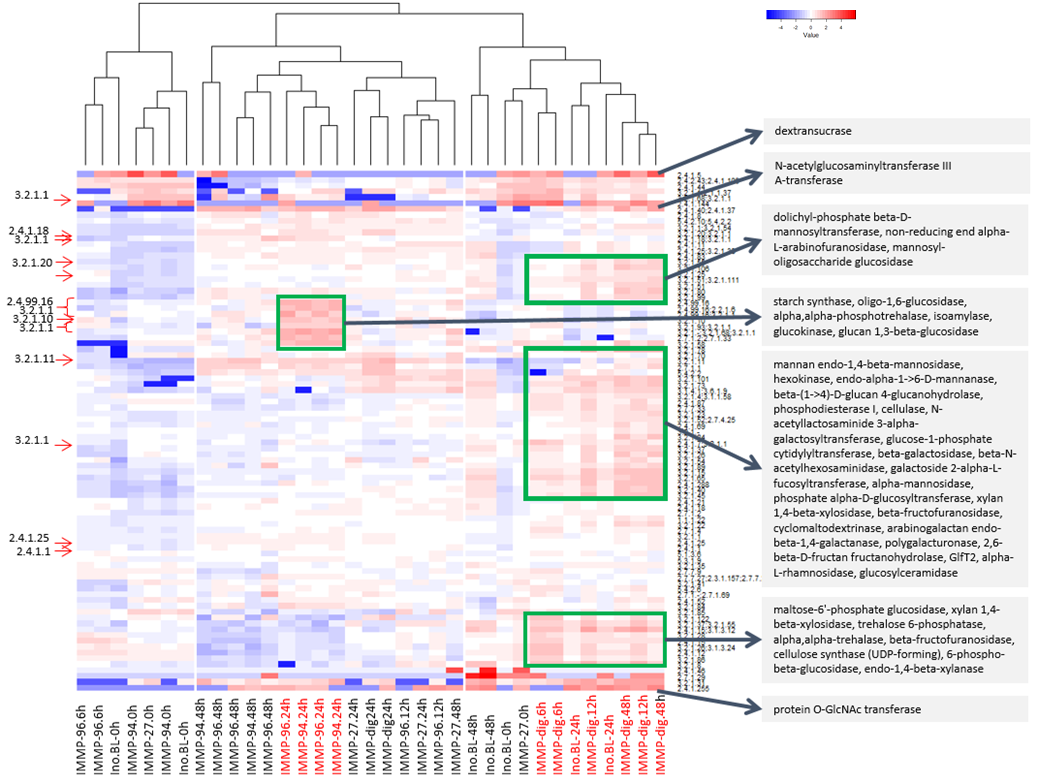

### Supplementary figure 6

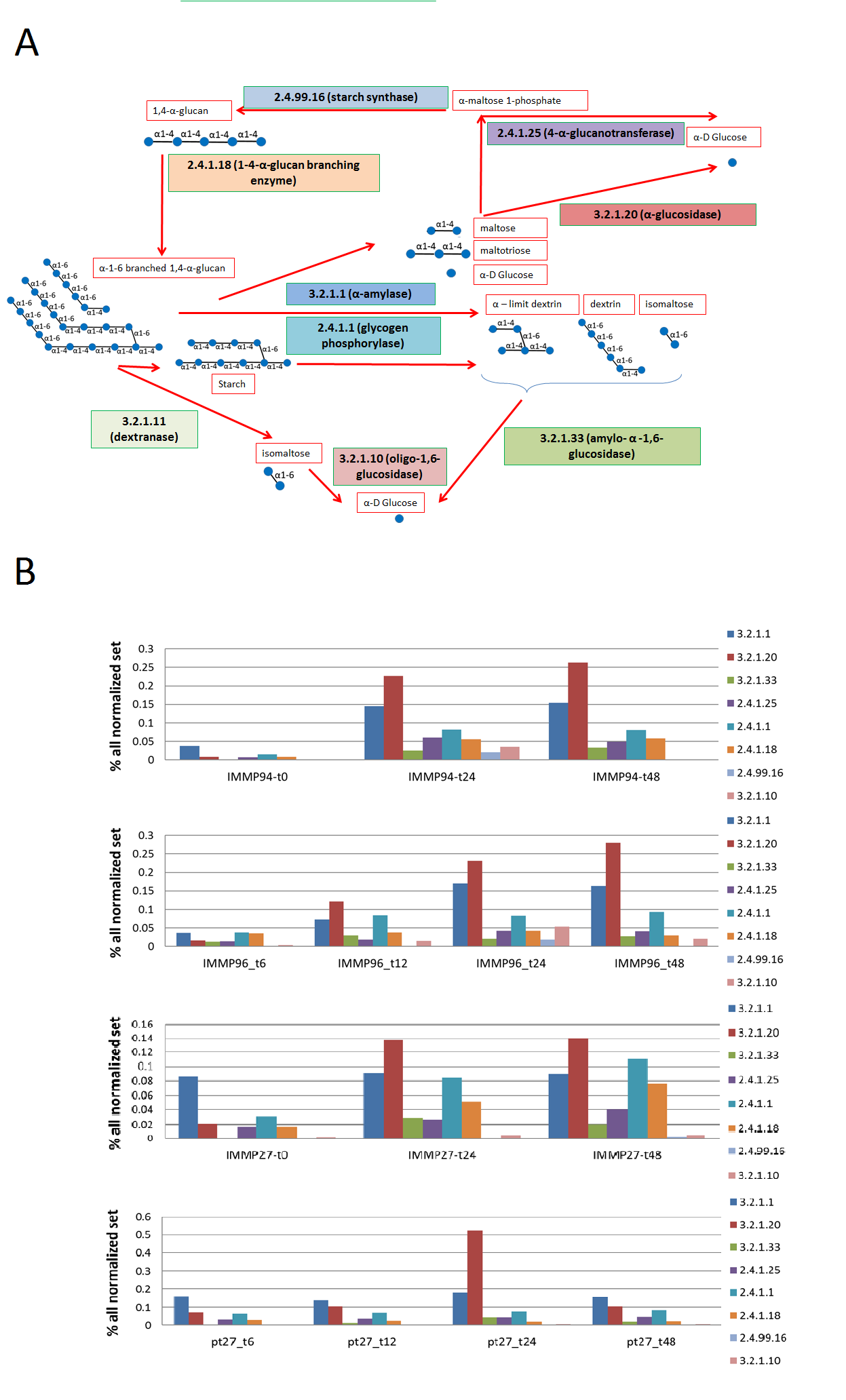

### Supplementary figure 7

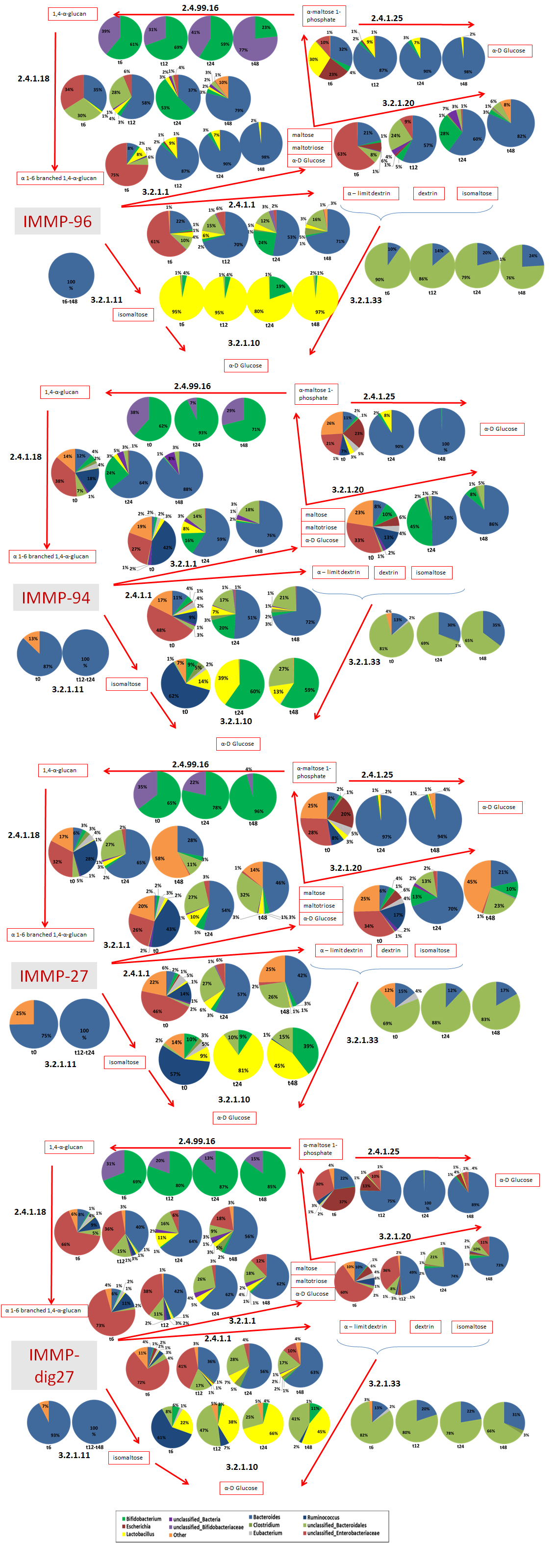
