## Supplementary materials and tables and figure legends for "Metatranscriptomic analysis indicates prebiotic effect of Isomalto/malto-polysaccharides on human colonic microbiota *in-vitro*"

Supplementary Material

Table S1. Overview over the RNA-seq metrics

| Condition | total reads | rRNA | % rRNA | non rRNA | trimmed bases due to adapters | % of bases trimmed due to adapters | sequences passing prinseq quality filtering | % passing prinseq quality filtering | mean length | Total % of bases passing ALL filtering steps | Mapping rate in % to assembly |
| --- | --- | --- | --- | --- | --- | --- | --- | --- | --- | --- | --- |
| Blank, repl. 1, t0 | 17182356 | 388147 | 2,26 | 16794209 | 713631983 | 28,14 | 15120013 | 90,03 | 108,34 | 63,56 | 79,06 |
| Blank, repl. 2, t0 | 12843968 | 234407 | 1,83 | 12609561 | 620076640 | 32,57 | 11339128 | 89,92 | 101,95 | 60,00 | 75,99 |
| IMMP-27, repl. 1, t0 | 18661592 | 402676 | 2,16 | 18258916 | 844672334 | 30,64 | 16454635 | 90,12 | 104,71 | 61,55 | 71,29 |
| IMMP-27, repl. 2, t0 | 21152485 | 438493 | 2,07 | 20713992 | 881059209 | 28,17 | 18728000 | 90,41 | 108,06 | 63,78 | 74,42 |
| IMMP-94, repl. 1, t0 | 30405866 | 555904 | 1,83 | 29849962 | 1087733238 | 24,13 | 26908763 | 90,15 | 113,98 | 67,25 | 79,06 |
| IMMP-94, repl. 2, t0 | 19354896 | 777385 | 4,02 | 18577511 | 588584413 | 20,98 | 16575459 | 89,22 | 119,53 | 68,24 | 80,79 |
| Blank, repl. 1, t24 | 27195831 | 243329 | 0,89 | 26952502 | 917731871 | 22,55 | 24835747 | 92,15 | 114,96 | 69,99 | 81,28 |
| Blank, repl. 2, t24 | 26279510 | 263802 | 1,00 | 26015708 | 1199020108 | 30,52 | 23724640 | 91,19 | 103,99 | 62,59 | 81,43 |
| IMMP-27, repl. 1, t24 | 12540536 | 1832716 | 14,61 | 10707820 | 587094636 | 36,31 | 9467522 | 88,42 | 93,59 | 47,10 | 85,32 |
| IMMP-27, repl. 2, t24 | 23402662 | 2094144 | 8,95 | 21308518 | 775851341 | 24,11 | 18915069 | 88,77 | 111,03 | 59,83 | 86,95 |
| IMMP-94, repl. 1, t24 | 14390413 | 901227 | 6,26 | 13489186 | 729841441 | 35,83 | 12186486 | 90,34 | 96,64 | 54,56 | 85,41 |
| IMMP-94, repl. 2, t24 | 27352727 | 1655263 | 6,05 | 25697464 | 1235521747 | 31,84 | 23084321 | 89,83 | 102,74 | 57,80 | 84,52 |
| Blank, repl. 1, t48 | 22532893 | 822320 | 3,65 | 21710573 | 1368385387 | 41,74 | 19533106 | 89,97 | 88,47 | 51,13 | 79,64 |
| Blank, repl. 2, t48 | 23432237 | 902055 | 3,85 | 22530182 | 1200860127 | 35,3 | 20328428 | 90,23 | 97,88 | 56,61 | 80,17 |
| IMMP-27, repl. 1, t48 | 24004294 | 4361225 | 18,17 | 19643069 | 1175519132 | 39,63 | 16574957 | 84,38 | 92,59 | 42,62 | 81,24 |
| IMMP-27, repl. 2, t48 | 21275156 | 8177863 | 38,44 | 13097293 | 691799504 | 34,98 | 11702497 | 89,35 | 98,7 | 36,19 | 80,22 |
| IMMP-96, repl. 1, t48 | 28735621 | 11137710 | 38,76 | 17597911 | 1296700269 | 48,8 | 14765189 | 83,9 | 81,2 | 27,82 | 83,92 |
| IMMP-96, repl. 2, t48 | 35679453 | 11638379 | 32,62 | 24041074 | 1511937150 | 41,65 | 20304145 | 84,46 | 92,59 | 35,13 | 85,36 |
| IMMP-96, repl. 1, t6 | 18066769 | 185180 | 1,03 | 17881589 | 390649819 | 14,47 | 15736624 | 88 | 126,02 | 73,18 | 75,76 |
| IMMP-96, repl. 2, t6 | 20993700 | 55456 | 0,26 | 20938244 | 412525567 | 13,05 | 18327046 | 87,53 | 128,66 | 74,88 | 71,55 |
| IMMP-96, repl. 1, t12 | 35944607 | 70088 | 0,20 | 35874519 | 1270799851 | 23,46 | 31773426 | 88,57 | 113,35 | 66,80 | 76,36 |
| IMMP-96, repl. 2, t12 | 22773537 | 64983 | 0,29 | 22708554 | 964291453 | 28,12 | 20307424 | 89,43 | 106,59 | 63,37 | 76,79 |
| IMMP-96, repl. 1, t24 | 17241186 | 1380292 | 8,01 | 15860894 | 385783659 | 16,11 | 13984773 | 88,17 | 122,24 | 66,10 | 87,1 |
| IMMP-96, repl. 2, t24 | 19319950 | 1502708 | 7,78 | 17817242 | 607265419 | 22,57 | 15673285 | 87,97 | 111,85 | 60,49 | 86,3 |
| IMMP-96, repl. 1, t48 | 34060532 | 145402 | 0,43 | 33915130 | 1420500568 | 27,74 | 30260173 | 89,22 | 107,64 | 63,75 | 75,81 |
| IMMP-96, repl. 2, t48 | 31992519 | 119756 | 0,37 | 31872763 | 1450909006 | 30,15 | 28116504 | 88,21 | 105,1 | 61,58 | 72,91 |
| Blank, repl. 1, t0 | 14780189 | 291372 | 1,97 | 14488817 | 458214833 | 20,94 | 12514201 | 86,37 | 111,32 | 62,84 | 84,1 |
| IMMP-dig27, repl. 1, t6 | 21177851 | 113633 | 0,54 | 21064218 | 1266290906 | 39,81 | 17927736 | 85,11 | 89,44 | 50,48 | 81,34 |
| IMMP-dig27, repl. 2, t6 | 14730499 | 91528 | 0,62 | 14638971 | 464265511 | 21 | 12827318 | 87,62 | 113,79 | 66,06 | 81,04 |
| IMMP-dig27, repl. 1, t12 | 19229982 | 120092 | 0,62 | 19109890 | 353792739 | 12,26 | 16872478 | 88,29 | 128,38 | 75,09 | 81,28 |
| IMMP-dig27, repl. 2, t12 | 19183847 | 184071 | 0,96 | 18999776 | 449569556 | 15,67 | 16837679 | 88,62 | 123,83 | 72,46 | 81,79 |
| IMMP-dig27, repl. 1, t24 | 19000262 | 346531 | 1,82 | 18653731 | 573909821 | 20,38 | 16612325 | 89,06 | 117,09 | 68,25 | 82,71 |
| IMMP-dig27, repl. 2, t24 | 18981915 | 260033 | 1,37 | 18721882 | 214922072 | 7,6 | 16320453 | 87,17 | 135,84 | 77,86 | 83,29 |
| IMMP-dig27, repl. 1, t48 | 19947312 | 1144473 | 5,74 | 18802839 | 508216740 | 17,9 | 16742425 | 89,04 | 120,37 | 67,35 | 88,6 |
| IMMP-dig27, repl. 2, t48 | 11159596 | 475984 | 4,27 | 10683612 | 160119185 | 9,93 | 9455765 | 88,51 | 132,49 | 74,84 | 89,42 |
| Average | 21366411,23 | 1514013,714 | 6,33 | 19852397,51 | 801840435,8 | 25,74 | 17591935,06 | 85,99 | 106,19 | 60,89 | 78,66 |
| Total | 747824393 | 52990480 |  | 694833913 | 28064415252 |  | 615717727 |  |  |  |  |

Table S3. Assignment of genus-level taxa per cluster, showing the amount of assigned genes and differentially expressed genes over all conditions

| **Organism** | **Genes assigned** | **Genes differentially expressed** |
| --- | --- | --- |
| **Cluster 1** |  |  |
| *Ruminococcus* | 9904 | 6220 |
| *Lactococcus* | 8211 | 5263 |
| unclassified*_*Gammaproteobacteria | 373 | 285 |
| Bos | 329 | 263 |
| N/A | 208 | 85 |
| unclassified_Mammalia | 198 | 67 |
| unclassified_Bovidae | 107 | 41 |
| Clostridiales | 4 | 0 |
| Bacteria | 3 | 1 |
| Eukaryota | 2 | 1 |
| Gammaproteobacteria | 1 | 1 |
| Viruses | 1 | 0 |
| **Cluster 2** |  |  |
| *Lactobacillus* | 8655 | 7279 |
| *Bifidobacterium* | 7817 | 5849 |
| unclassified_Actinobacteria | 265 | 195 |
| unclassified_Bifidobacteriaceae | 116 | 98 |
| *Fusobacterium* | 70 | 47 |
| unclassified_Lactobacillaceae | 52 | 38 |
| Myoviridae | 36 | 33 |
| **Cluster 3** |  |  |
| *Enterococcus* | 2580 | 1927 |
| unclassified_Lactobacillales | 1154 | 1012 |
| unclassified_Bacilli | 537 | 473 |
| unclassified_Enterococcaceae | 139 | 136 |
| **Cluster 4** |  |  |
| unclassified_Enterobacteriaceae | 11350 | 9255 |
| unclassified_Bacteria | 6997 | 4494 |
| *Eubacterium* | 4884 | 3849 |
| *Escherichia* | 2972 | 2386 |
| unclassified*_*Proteobacteria | 683 | 530 |
| *Salmonella* | 59 | 47 |
| *Shigella* | 51 | 9 |
| *Enterobacter* | 36 | 29 |
| *Citrobacter* | 31 | 14 |
| *Vibrio* | 26 | 9 |
| **Cluster 5** |  |  |
| *Bacteroides* | 53749 | 49101 |
| unclassified*_*Bacteroidales | 10243 | 9067 |
| *Parabacteroides* | 1749 | 919 |
| *Prevotella* | 302 | 233 |
| *Bacteria* | 193 | 133 |
| *Desulfosporosinus* | 32 | 11 |
| *Flavobacterium* | 30 | 27 |
| **Cluster 6** |  |  |
| *Clostridium* | 4244 | 3228 |
| unclassified*_*Clostridia | 114 | 69 |
| **Cluster 7** |  |  |
| unclassified_Clostridiales | 10819 | 3356 |
| unclassified_Lachnospiraceae | 1030 | 219 |
| *Anaerostipes* | 127 | 6 |
| Clostridiales | 39 | 5 |
| **Cluster 8** |  |  |
| N/A | 8225 | 4083 |
| *Sutterella* | 3598 | 2645 |
| unclassified_Bacteroidetes | 770 | 681 |
| unclassified_Betaproteobacteria | 135 | 77 |
| *Odoribacter* | 94 | 76 |
| *Ethanoligenens* | 28 | 21 |
| *Corynebacterium* | 20 | 8 |
| **Cluster 9** |  |  |
| *Bilophila* | 3853 | 2614 |
| unclassified*_*Firmicutes | 3152 | 1898 |
| *Phascolarctobacterium* | 1328 | 8 |
| unclassified*_*Selenomonadales | 244 | 2 |
| *Acidaminococcus* | 174 | 4 |
| unclassified*_*Acidaminococcaceae | 117 | 0 |
| *Selenomonas* | 109 | 19 |
| *Veillonella* | 88 | 8 |
| *Megamonas* | 64 | 12 |
| unclassified*_*Veillonellaceae | 56 | 3 |
| *Pelosinus* | 53 | 4 |
| *Megasphaera* | 53 | 9 |
| *Desulfitobacterium* | 51 | 15 |
| *Anaeromusa* | 33 | 0 |
| *Acetonema* | 32 | 3 |
| *Mitsuokella* | 20 | 5 |

**Supplementary Methods**

Text mining

Further EC numbers were derived by text mining and matching all InterProScan derived domain names against the BRENDA database (download 13.06.13) [23]. The text mining algorithm included lower casing all characters, removal of non-alphanumerical characters (colons, commas, brackets, apostrophes, dashes, terminal points), removal of partial and generic terms (type, terminal, subunit, domain, enzyme, like, hypothetical, conserved, operon, active site, enzyme, probably, central, 51 kd, respiratory chain, c terminal, n terminal), rejection of overly generic final result terms (kinase, cytochrome, protein, methyltransferase) and reduction of certain terms (deletion of PEP/pyruvate binding; removal of “prokaryotic” in “prokaryotic cytidylate kinase”; “family” in “cytidilate kinase family”; “phosphorylating” in “glyceraldehyde phosphate dehydrogenase phosphorylating”; “iron containing” in “iron containing alcohol dehydrogenase”; “zinc containing” in “zinc containing alcohol dehydrogenase”; “manganese containing” in “manganese containing catalase”; “20 kd” in “nadh ubiquinone oxidoreductase 20 kd”; replacement of “carboxyltransferase” with “carboxylase” in “pyruvate carboxyltransferase”). Furthermore, all terms, which were only of length one, were also removed, in case the remaining name contained more than two words. On some domain names a manual curation was performed, and overly generic identifications (e.g. matching PF12847 “Methyltransferase domain” with e.g. EC 2.1.1.124 with alternative name “Protein Methyltransferase I”) were rejected.

**Supplementary Figures**

Figure S1: Experimental design. We would like to thank Bartholomeus van den Bogert for the design of this figure.

Figure S2: Overview over the clustering procedure. First, expression was lumped at the genus level. On the accumulated expression data k-means clustering was performed, until a stable clustering was achieved. The genes of the grouped genera were afterwards subjected to DBSCAN clustering. The stability of the clustering was evaluated with the Tau-parameter. Only genes, which were at least once differentially expressed, were used in the clustering process to reduce the noise.

Figure S3. Overview of the main gene expression patterns. All groups (*Bacteroides, E.coli, Lactobacillus/Bifidobacterium, Enterococcus*; besides *Eubacterium hallii*) showed in all prebiotic conditions increase in relative transcript abundance in roughly the same proportion (green). Some groups (*Bacteroides, Escherichia*) also showed comparable increase in expression in the control condition (dotted green line). Furthermore, all groups showed a downregulation of certain genes in all conditions (red), and an upregulation of a group of genes in the control condition (black). *Eubacterium hallii* showed only increase in transcript abundance at the last time point with the prebiotic IMMP-27 (yellow).

Figure S4. Relative abundance (percentage) of starch and sucrose metabolism enzyme encoding genes detected in the metatranscriptome data.

Figure S5. Heatmap of log10 transformed relative abundances of expressed genes detected in our data coding for starch and sucrose metabolism enzymes. Samples clustered based on the similarities between the up and down regulated genes. The red arrows indicate selected genes that code for enzymes described in our IMMP degradation model. Green boxes highlight the gene upregulation patterns for different IMMPs at various incubation times.

Figure S6. IMMP degradation model. a. main enzymes involved in the pathway, b. relative abundance of transcripts of genes coding for enzymes needed for IMMP degradation.

Figure S7. Relative contribution of different bacterial groups to expression of genes coding for the enzymes in the IMMP degradation pathway.
